# Spatiotemporal properties of common semantic categories for words and pictures

**DOI:** 10.1101/2023.09.21.558770

**Authors:** Yulia Bezsudnova, Andrew J. Quinn, Ole Jensen

**Affiliations:** Centre for Human Brain Health, School of Physics and Astronomy, University of Birmingham, UK; Centre for Human Brain Health, School of Psychology, University of Birmingham, UK

## Abstract

The timing of semantic processing during object recognition in the brain is a topic of ongoing discussion. One way of addressing this question is by applying multivariate pattern analysis (MVPA) to human electrophysiological responses to object images of different semantic categories. However, while MVPA can reveal whether neuronal activity patterns are distinct for different stimulus categories, concerns remain on whether low-level visual features also contribute to the classification results. To circumvent this issue, we applied a cross-decoding approach to magnetoencephalography (MEG) data from stimuli from two different modalities: images and their corresponding written words. We employed items for three categories and presented them in a randomized order. We show that If the classifier is trained on words, pictures are classified between 150 - 430 ms from after stimulus onset, and when training on pictures, words are classified between 225 - 430 ms. The topographical map identified using a searchlight approach for cross-modal activation in both directions showed left lateralization confirming the involvement of linguistics representations. These results point to semantic activation of pictorial stimuli occurring at ≈150 ms whereas for words the semantic activation occurs at ≈230 ms.

## 1 Introduction

Humans are capable of recognizing and inferring the semantic category of a presented object regardless of the modality in which it is presented, whether through visual, auditory, or textual means.

A powerful way of studying how the human brain encodes the semantics of objects is to apply multivariate pattern analysis (MVPA) to human electrophysiological responses to stimuli of different conceptual categories (Cichy et al., 2014; Groen et al., 2022; Carlson et al., 2013). The timing of the transition from visual representations of individual objects to more abstract semantic-type concepts is determined by identifying the time points at which the category of the object can be accurately predicted from the multivariate representation in the MEG and EEG data (Proklova et al., 2016; Kumar et al., 2017). However, drawing meaningful conclusions from the above-chance classification using stimuli from a single modality may be limited, as there are often multiple dimensions in which the two conditions differ (Peelen and Downing, 2022; Frisby et al., 2023). For example, one can argue that perceptual information beyond semantic content influences classification outcomes. Furthermore, the temporal dynamic of object recognition may also be influenced by the specific experimental task employed in the study, such as a one-back task, category judgment, or detection task. As such the precise timing of semantic category activation remains an open question (Giari et al., 2020; Kaiser et al., 2016; Dirani and Pylkkänen, 2023; Miozzo et al., 2015; MacGregor et al., 2012; Dirani and Pylkkänen, 2020; Simanova et al., 2010).

Studying brain responses to stimuli from different modalities (e.g. words versus images) mitigates the criticism related to the influence of perceptual features on classification, as these stimuli do not share common low-level features (Hauk et al., 2006). Additionally, demonstrating the existence of cross-modal generalization (training a classifier on one modality, classifying another) at specific time points will provide insights into the ongoing debate about how and when semantic information is encoded. This includes testing the hub-and-spoke theory of semantic representations where the representation first arises in a modality-specific way and later transfers to the amodal hub (Ralph et al., 2017; Humphreys et al., 2015; Pobric et al., 2007). There is limited research done to investigate the time of cross-modal generalization using M/EEG and the reported time dynamic of cross-modal representation varies (Leonardelli et al., 2019; Dirani and Pylkkänen, 2023; Iamshchinina et al., 2022; Simanova et al., 2010).

In the MEG study (Leonardelli et al., 2019) the categorization between famous places (“Big Ben”) and famous people (“Brad Pitt”) is studied for the picture and written word modalities. After each stimulus, participants were asked to perform either a shallow categorization task (Place or Person?) or a deeper semantic task (“Italian or foreign?”). Within the modality (modality-specific) category information robustly appears around 100 ms for pictures and 230 ms for words. The cross-modal (across modality) classification was studied only from words to pictures and revealed three significant clusters separated in time. Specifically, the authors suggest that cross-modal generalization unfolds through a three-stage process and the first shared representations are accessed at 200 ms using words and at 110 ms using pictures. However, brain activity from concepts such as famous places and people used in their paradigm might not generalize to more common objects. Furthermore, any findings of cross-modal representations across modalities could be driven by a shared category-judgment process due to a category-naming task rather than an automatic activation of semantic representations (Hebart et al., 2018).

Another recent MEG study (Dirani and Pylkkänen, 2023) examining generalization between written words and pictures using picture naming and word reading tasks showed different results. Significant decoding (animal vs. tool) activates surprisingly early around 75 ms for pictures, and 95 ms for words. Cross-generalization occurs simultaneously for both modalities around 150 ms. The different time courses of the shared semantic representation between (Dirani and Pylkkänen, 2023) study and (Leonardelli et al., 2019) study could be attributed to variations in participant tasks and experimental paradigms. Importantly, in (Dirani and Pylkkänen, 2023) a block presentation of categories was utilized to demonstrate cross-modal categorization. This might result in anticipatory effects making the semantic categorization occur earlier. Based on these considerations, the precise timing of the activation of the shared representation should be reevaluated in experiments where the category order is randomized while explicit category naming is not required.

In this work, we investigated the time course of semantic activation within and across pictural and textual stimulation presented in a randomized order. We analyzed MEG data using MVPA analysis. To further eliminate any anticipatory biases such as the possibility of stronger activation of general motor commands for the tools category we chose to include multiple categories. Finally, we applied a searchlight approach to identify the brain areas involved in the semantic representations.

## 2 Methods

### 2.1 Participants

Thirty-eight healthy adult participants took part in the study. Five participants had to be excluded due to extensive noise in the data and two participants were excluded due to low accuracy during the task (accuracy less than 85%). Therefore the final sample consisted of 31 participants (mean age *pm* SD = 21.77 *pm* 3.31; 21 female). The number of participants was selected to match the average number of participants used in previous studies that employed MVPA analysis (Leonardelli et al., 2019; Dirani and Pylkkänen, 2023; Singer et al., 2023). The study was conducted at the Center of Human Brain Health in Birmingham, United Kingdom. All participants were native English speakers with normal or corrected-to-normal vision. The University of Birmingham Ethics Committee approved the study. The participants provided written informed consent and received £15 per hour or course credits as compensation for their participation.

### 2.2 Experimental paradigm

The experimental design was inspired by the study (Iamshchinina et al., 2022), however, some stimuli were replaced as well as the task for the participant was modified. The stimulus set was composed of 48 objects. Each stimulus was presented as a picture and a written word. The objects were organized according to three dimensions of categories, each separated into two categorical divisions: “size” (big or small), “movement” (moving or still), and “nature” (natural or man-made). Each object in the study belonged to each of the dimensions (e.g. a book is small, still, and man-made). The stimulus set was balanced such that each categorical division included one-half of the stimulus set (24 objects). Hence each set of categories (e.g. big, still, natural) included 6 objects. The selection of categorical divisions in this study was based on previous findings demonstrating reliable neural representations of semantic dimensions across categories (Iamshchinina et al., 2022; Konkle and Caramazza, 2013). For other purposes not to be analyzed here, each object is presented in two modalities: written words and pictures. The size of the images is 400 x 400 pixels presented on a grey screen at a visual angle of 6 °. In the stimuli set for textual modality (font: Arial, bold, 75), we did not control for the lexical properties of the words, which resulted in a mixture of high-frequency and low-frequency words. Additionally, the length of the words was not controlled.

Each object (word or picture) was presented for 0.6 s. Prior to the object presentation, the fixation cross was shown for 0.5-0.7 s. The experiment was divided into 9 blocks where either the succession of words (5 blocks) or images (4 blocks) was presented. The experimental design followed a consistent order, with word blocks always presented first, followed by image blocks, and so on (Fig 1B). After each block, a participant had a break. We included an extra word block based on previous reports that indicated a lower signal-to-noise ratio for classification in the textual modality (Simanova et al., 2010; Dirani and Pylkkänen, 2023). Each block consisted of 240 trials, with 48 of them being question trials, and lasted for around 6 minutes. Each stimulus was present 4 times in a block: when pictures were presented the probe question was presented as a word and vice versa (Fig 1B). As such a participant had to identify the correct picture during the written words and vice versa. This design aimed to keep participants engaged and stimulate deep cross-modal perception without explicitly addressing the category distinction between stimuli. Note that in each block the probe question was applied for every object once and alternative choices were unique as well.

**Figure 1:**
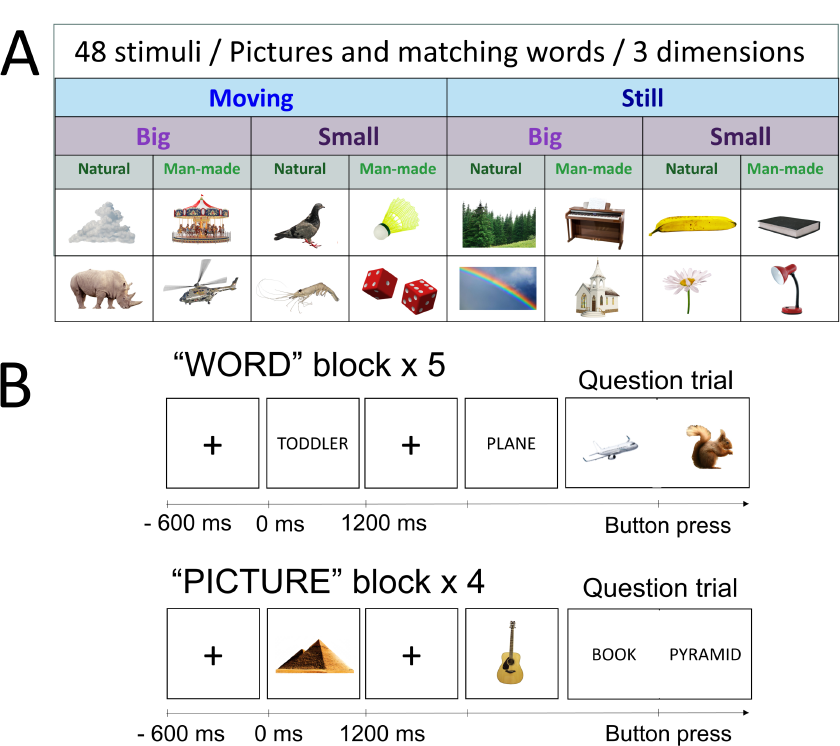
Experimental paradigm. A) The stimulus set is comprised of 48 objects which can be divided according to 3 dimensions (“size”: big/small; “movement”: still/moving; “nature”: natural/man-made). B) In both image and word blocks, participants were presented with stimuli in random order. In the picture block, participants viewed images of the objects, while in the word block, they read the corresponding words. The task (question trial) was presented randomly every fifth trial on average. Participants were required to identify whether the picture or word corresponded to a previously seen stimulus and press the appropriate button accordingly. When pictures were presented the probe question was presented as a word and vice versa.

The stimuli presentation is implemented in Matlab using the Psychophysics Toolbox (Brainard and Vision, 1997).

### 2.3 MEG acquisition

MEG data were recorded using a 306-sensor TRIUX MEGIN Elekta Neuromag system, consisting of 204 orthogonal gradiometers and 102 magnetometers, with online band-pass filtering from 0.1 to 330 Hz and a sampling rate of 1000 Hz. Before collecting data, the positions of three anatomical fiducial points (nasion, left preauricular, and right preauricular points) were recorded using a Polhemus Fastrack electromagnetic digitizer system. Furthermore, we recorded the positions of four head-position indicator coils (HPI coils): two placed on the left and right mastoid bone, and two on the forehead, with a minimum separation of 3 cm between each coil. After completing these initial steps, participants were seated in an upright position with a 60° angled backrest within the MEG gantry. Electrodes were affixed about 2.5 cm from the outer canthus of each eye to capture horizontal EOG signals. To record the vertical EOG, a pair of electrodes was placed above and below the right eye in line with the pupil. The ECG signals were captured using a set of electrodes positioned on both the left and right collarbones.

### 2.4 Data preprocessing

The data are analyzed using the open-source toolbox MNE Python v1.4.2 (Gramfort et al., 2013) following the standards defined in the FLUX Pipeline (Ferrante et al., 2022).

First, blinks and muscle artefacts are annotated using EOG channels and magnetometers recordings respectively. A semi-automatic detection algorithm was utilized to mark sensors with excessive artefacts (on average 4.5 channels per participant). Then, the data was low-pass filtered at 100Hz to reduce HPI coils artefacts. We did not use signal-space separation (SSS) or Maxwell filtering because our previous research demonstrated that they negatively impact classification results (in preparation). We attribute this decline in performance to the significant rank reduction resulting from SSS filtering (Taulu and Simola, 2006). Independent Component Analysis (ICA) algorithm was applied (Hyvärinen and Oja, 2000) to remove components associated with cardiac artefacts and eyeblinks. The identification process involved analyzing the time courses and topographies of the ICA components. On average, 3 components corresponding to two cardiac-related and one blink-related artefact were identified and removed for each subject. After ICA the data was segmented into trials. Trials corresponding to the same stimulus were averaged together to construct “supertrials”. This step further enhances the signal-to-noise ratio (SNR) of the data (Ashton et al., 2022; Guggenmos et al., 2018). We applied a highpass filter of 0.1 Hz and downsampled the data to 500 Hz. The epochs were time-locked to the onset of the stimuli and were cropped to a time window of 100 ms before the stimuli and 700 ms after the stimuli onset. Each time point was represented as a 306-dimensional vector (channel data). We expanded this vector by including past 25 ms (12 time points) and future 25 ms (12 time points) of data, resulting in a 306×25 dimensional feature vector representing each time point. This procedure (termed “delayed embedding”) resulted in a more information-rich representation of the neural activity associated with a given stimulus by incorporating more time points (Chan et al., 2011; Tyler et al., 2013; Cheng, 2021). This feature vector was then used in the classifier.

### 2.5 Classification analysis

Multivariate pattern analysis was applied to the pre-processed MEG data to classify the categories of the presented objects (Haynes and Rees, 2006; Cichy et al., 2014; Carlson et al., 2013). Trials associated with different categories (e.g. moving vs still) and one modality (e.g. words) are labelled accordingly and used as input for the classifier. Prior to classification, the data was standardized by removing the mean and scaling to unit variance per sensor. Subsequently, we employed a support vector machine (SVM) (Cortes and Vapnik, 1995) from the Python module Scikit-learn to classify the data over time. The classification procedure relied on a 5-fold cross-validation approach, and the performance was quantified using the area under the curve (AUC) metric, which measures the classifier discriminative ability. This procedure was done separately for each subject.

First, we investigated the classification accuracy with time for each category dimension (“movement”, “size”, and “nature”) averaged across participants for each modality separately. The dimensions where both modalities showed classification accuracy significantly above the chance level were selected for cross-modal analysis. For cross-modal analysis, we also used time-generalized MVPA (King and Dehaene, 2014). We used the classification approach mentioned above with the exception that the classifier was training in one modality at one time point and testing it on another at a different time point. This procedure was done twice, once with the classifier trained on the words data and tested on the pictures data, and once where it was trained on the pictures data and tested on the words.

In order to explore the spatial distribution of representations across the MEG sensors over time, we employed the searchlight approach on sensor-level data (Leonardelli et al., 2019; Kriegeskorte et al., 2008). In this approach, classification was performed on “patches” which are defined by one sensor (e.g. sensor MEG 1423) and all sensors within a 4 cm radius (e.g. 4 cm from MEG 1423). This typically resulted in 15 sensors (consisting of gradiometers and magnetometers). Therefore the feature vector had dimensions Nx25, where N is the number of sensors in the “patch” and 25 time points from the delayed-embedding procedure. Patches were created for all possible sensor locations (rim sensors had fewer sensors in the patch). For each channel location the classification accuracy from the relevant “patch” was averaged over a chosen time interval and this value was plotted on the topographical sensor map.

### 2.6 Statistical analysis

To find significant time points for the modality-specific classification curves and simultaneous cross-generalization, we used a non-parametric one-sampled permutation t-test (one-tailed) against 50% chance level, controlled for multiple comparisons (Sassenhagen and Draschkow, 2019; Maris and Oostenveld, 2007) implemented in the GLMTools Python package (https://pypi.org/project/glmtools/). We set a cluster-forming threshold of 1.7, corresponding to an alpha threshold of 0.05. Clusters of t-values that exceeded the cluster-forming threshold were formed based on direct adjacency in time (the minimum number of vertices in terms of time points in a cluster was set to 2) and summarised using the sum of t-values within the cluster. The largest cluster from each of N sign-flip permutations were computed to form a null distribution. A cluster in the observed t-values was considered significant if its cluster stat (sum of t-value with cluster) lay at or above the 95th percentile of the null distribution. This corresponds to an alpha threshold of p=0.05.

The same analysis was done for cross-modal time generalization results. Note that in this case the clusters are based on direct adjacency in both axes.

## 3 Results

### 3.1 Within-modality decoding

The category decoding shows significant results for pictures for every categorical dimension (“size”, “movement”, and “nature”) separately. For words, we found the above channel level classification only for dimensions “size” (big/small) and “movement” (still/moving), but not “nature” (natural/man-made). In a previous study (Cichy et al., 2014) this category also exhibited the lowest decoding accuracy when examining pictures, suggesting that it might also be difficult to decode across modalities (Dirani and Pylkkänen, 2023; Iamshchinina et al., 2022). Therefore, all the results were derived by averaging the classification results obtained from just two categorical dimensions (“size” and “movement”).

The classification accuracies for the picture modality and textual modality are shown in Fig. 2A. As expected, a classifier that uses brain activity elicited by words showed lower performance compared to picture stimuli. For words, decoding is most pronounced around 240 - 350 ms after stimuli are presented. For pictures, decoding is most pronounced around 155 - 510 ms. These results indicate that semantic activation for words occurs approximately 100 ms later than for pictures. Note that the time estimation might be blurred due to the feature vector including ±25 ms of information, however, the relative relationships between modalities are preserved.

**Figure 2:**
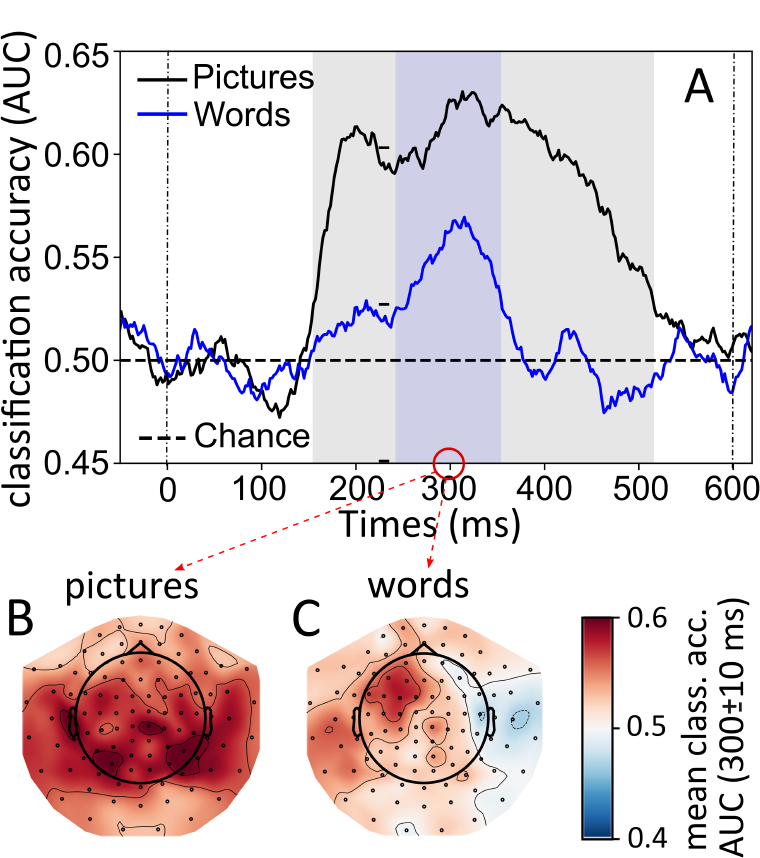
A) The time course of category decoding averaged over two categorical dimensions: “size” (big/small), and “movement” (still/moving). The blue line marks the classification curve for word modality, black line marks the classification curve for picture modality. Significant clusters (p *<* 0.05; controlled for multiple comparisons over time) are shown as highlighted areas accordingly; B) Topographical map of category decoding averaged over 300±10 ms created using searchlight MVPA decoding categories of pictures, C) decoding categories of words. The colourcode indicates the classification accuracy (AUC) averaged over the time interval 300±10 ms. The classification accuracy is averaged over two category dimensions: “size” (big/small), “movement” (still/moving)

For the time points (300±10 ms) when the decoding accuracy is strongest, we show topographical maps of the classification accuracy using a searchlight approach. The sensors that contributed strongest to the overall accuracy of the picture category are located over bilateral temporal and parietal areas (Fig. 2B). For the decoding word category, the informative sensors are located over the left temporal and left frontal parts of the brain (Fig. 2C). This localization is in line with prior results using MEG in object decoding (Cichy et al., 2014) and word reading studies (Kumar et al., 2017; Santi et al., 2015).

### 3.2 Cross-modal decoding

The simultaneous (trained and tested on the same time points) cross-modal classification between textual and pictural modality is shown in Fig. 3A. When training on words and testing on pictures, the significant cluster emerged at 280 to 430 ms. When reversely training on pictures and testing on words, the cluster emerged at 330 to 430 ms.

**Figure 3:**
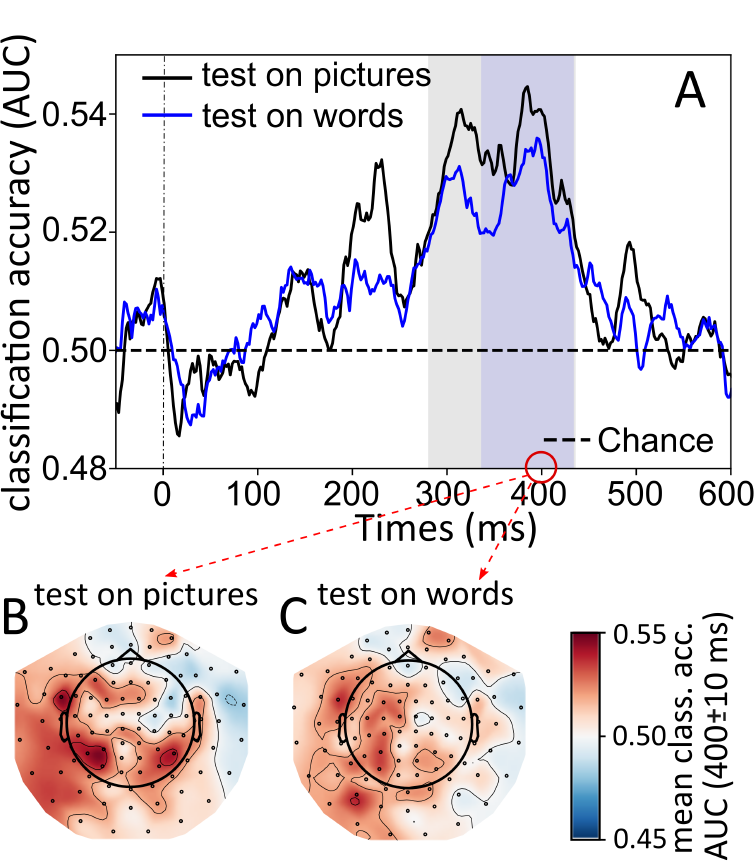
A) The time course of simultaneous (trained and tested on the same time points) cross-modal category decoding averaged over two categorical dimensions: “size” (big/small), and “movement” (still/moving). The blue line marks the classification curve when classifiers were trained on pictures and tested on words, black line marks the classification curve when trained on words and tested on pictures. Significant clusters (p *<* 0.05; controlled for multiple comparisons over time) are shown as highlighted areas accordingly; B) Topographical map of cross-modal categorical decoding averaged over 400±10 ms created using searchlight MVPA: when training on words, testing on pictures modality; C) when training on pictures, testing on words modality. The colourcode indicates the classification accuracy averaged over the time interval 400±10 ms. The classification accuracy is averaged over two categorical dimensions: “size” (big/small) and “movement” (still/moving).

Next, we show the topographical maps of the decoding accuracy at 400±10 ms to check where the simultaneous cross-modal classification is the most pronounced (Fig. 3B,C). We selected the time interval of 400±10 ms because it corresponds to the peak decoding accuracy in Fig. 3A. These maps for simultaneous cross-modal activation in both directions show more pronounced left lateralization (Fig. 3B,C) compared to topographies from within-modality decoding (Fig. 2B,C).

We further examined time-generalized cross-decoding results.

When training on pictures and testing on words the shared representation occurs between 225 - 430 ms (Fig. 4A); when training on words and testing on pictures the shared representation occurs between 150 and 430 ms (Fig. 4B). Interestingly, the most pronounced semantic decoding across modalities for both words and pictures becomes significant around the same time as the categories are classified within each modality (Fig. 2A). In sum, when considering the within- and cross-modal results, this point to semantic activation of pictures and words occurs at 150 ms and 230 ms respectively.

**Figure 4:**
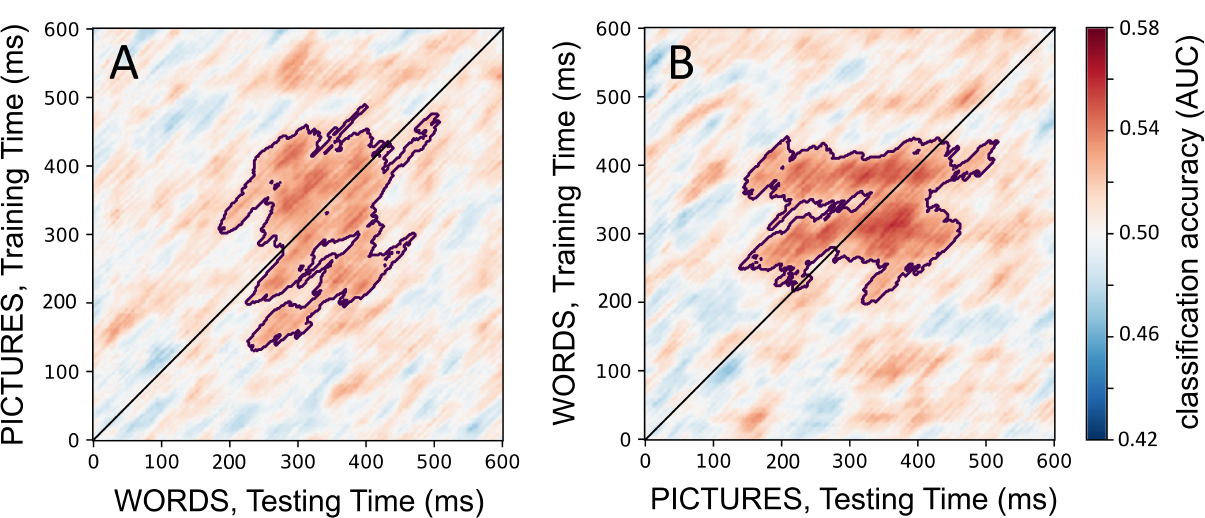
Time generalization results for category information, where classifier was A) trained on pictures and tested on words; B) trained on words and tested on pictures. The classification accuracy is averaged over two categorical dimensions: “size” (big/small), and “movement” (still/moving). Significant clusters are highlighted (one-sided permutation test, p ¡ 0.05, corrected for multiple comparisons).

## 4 Discussion

In this study, we investigated the temporal dynamics of semantic processing when objects were as pictures and text. We assessed the qualitative similarities between categories elicited for each modality using MVPA analysis on MEG data. The classification shows the most pronounced decoding activation for the pictures starting at 155 ms over the bilateral posterior part of the brain and for words after ≈240 ms over the left temporal and frontal parts of the brain (Fig. 2). We show successful cross-modal classification using MEG data (Fig. 4). If the classifier is trained on words, pictures are classified between 150 - 430 ms, and when training on pictures, words are classified between 225 - 430 ms. The topographical map for simultaneous cross-modal activation in both directions (from words to pictures and from pictures to words) reveals strong left lateralization (Fig. 3B,C).

Semantic activation for pictures starts around 150 ms, consistent with findings reported in a previous study (Iamshchinina et al., 2022) that employed a similar set of stimuli. The temporal dynamics of representations elicited by words demonstrate a later activation around 240 ms, which is in line with (Leonardelli et al., 2019) and studies using the N400 paradigm where the N400 response starts to build up at 250 ms (Federmeier, 2022). The different timing of picture and word categorization can be explained by the fact that low-level visual features in written words are less informative of semantic categories compared to pictures (Hauk et al., 2006). Note that decoding onsets of 75 ms for pictures and 95 ms for words acquired in the study (Dirani and Pylkkänen, 2023) is almost 70 ms earlier than what we have demonstrated here. This disparity could likely be attributed to the categorical presentation in blocks as in the study by Dirani et al. (Dirani and Pylkkänen, 2023), which facilitates faster category extraction.

The time course of cross-modal decoding (Fig. 4) with off-diagonal time points structured as a rectangle suggests the shared semantic representation exhibits some sustained features common to both modalities. Semantic activation when classifying from pictures to words starts around 225 ms, while from words to pictures, it begins approximately at 150 ms. We attribute comparable timing of modality-specific and cross-modal semantic decoding to linguistic activation when viewing both pictures and words due to the task design. Participants had to identify the word that corresponds to the previously seen picture and vice versa, therefore the association between the two modalities was encouraged. This also explains why the topographical maps from cross-modal decoding for both directions (Fig. 3B,C) show that the most informative sensors are located over the left regions of the brain, resembling the topographical map of word categorization. Words elicit abstract shared category representations due to their non-representative low-level feature, whereas categorization within pictorial modality is contaminated by perceptual features. Furthermore, the location of most informative sensors may include the left anterior temporal lobe (ATL) as shown with fMRI to be involved in semantic processing across various tasks and, including cross-modal generalization between written words and corresponding images (Humphreys et al., 2015; Branzi et al., 2020). These studies support the hub and spoke theory (Ralph et al., 2017) in which the ATL supports shared representations. While our MEG study is inconsistent with the hub and spoke theory for semantic representations, our topographical plots do point to an extended network supporting the semantic encoding going beyond the ATL.

## 5 Conclusion

Our results demonstrate the time course of modality-independent semantic representations isolated from perceptual confounds. Specifically, we examined the time course of activation of semantic representations common for words and pictures. We found that the semantic activation of words occurs at ≈230 ms. In the future, we will conduct the same experiment with OPM-MEG system to check our hypothesis that the new system can offer significant advantages in experiments designed for multivariate pattern analysis.

## 6 Data availability

Raw data is available upon request and code can be accessed on https://github.com/Y-Bezs/cross-modal-project

## 7 Authors contribution

Yulia Bezsudnova: Conceptualization; Formal analysis; Investigation; Methodology; Project administration; Visualization; Writing—Original draft; Writing—Review & editing. Andrew Quinn: Statistical analysis. Ole Jensen: Conceptualization; Supervision; Funding Acquisition; Writing—Review & editing.

## 8 Acknowledgements

The computations described in this paper were performed using the University of Birmingham’s BEAR Cloud service, which provides flexible resources for intensive computational work to the University’s research community. See http://www.birmingham.ac.uk/bear for more details. We thank Jonathan Winter for his support of MEG data acquisition. We also thank Yali Pan, and Syanah Wynn for the fruitful discussions.

## 9 Funding information

This work was supported by the BBSRC (grant number BB/R018723/1), and an EPSRC (grant number EP/T001046/1). And a Wellcome Trust Investigator Award in Science (grant number 207550).

## Notes

### Competing Interest Statement

The authors have declared no competing interest.

## References

Ashton, K., Zinszer, B. D., Cichy, R. M., Nelson III, C. A., Aslin, R. N., and Bayet, L. (2022). Time-resolved multivariate pattern analysis of infant eeg data: A practical tutorial. Developmental cognitive neuroscience, 54:101094.

Brainard, D. H. and Vision, S. (1997). The psychophysics toolbox. Spatial vision, 10(4):433–436.

Branzi, F. M., Humphreys, G. F., Hoffman, P., and Ralph, M. A. L. (2020). Revealing the neural networks that extract conceptual gestalts from continuously evolving or changing semantic contexts. NeuroImage, 220:116802.

Carlson, T., Tovar, D. A., Alink, A., and Kriegeskorte, N. (2013). Representational dynamics of object vision: the first 1000 ms. Journal of vision, 13(10):1–1.

Chan, A. M., Halgren, E., Marinkovic, K., and Cash, S. S. (2011). Decoding word and category-specific spatiotemporal representations from meg and eeg. Neuroimage, 54(4):3028–3039.

Cheng, F. (2021). Using single-trial representational similarity analysis with eeg to track semantic similarity in emotional word processing. arXiv preprint arXiv:2110.03529.

Cichy, R. M., Pantazis, D., and Oliva, A. (2014). Resolving human object recognition in space and time. Nature neuroscience, 17(3):455–462.

Cortes, C. and Vapnik, V. (1995). Support-vector networks. Machine learning, 20:273–297.

Dirani, J. and Pylkkänen, L. (2020). Lexical access in naming and reading: spatiotemporal localization of semantic facilitation and interference using meg. Neurobiology of Language, 1(2):185–207.

Dirani, J. and Pylkkänen, L. (2023). The time course of cross-modal representations of conceptual categories. NeuroImage, page 120254.

Federmeier, K. D. (2022). Connecting and considering: Electrophysiology provides insights into comprehension. Psychophysiology, 59(1):e13940.

Ferrante, O., Liu, L., Minarik, T., Gorska, U., Ghafari, T., Luo, H., and Jensen, O. (2022). Flux: A pipeline for meg analysis. NeuroImage, 253:119047.

Frisby, S. L., Halai, A. D., Cox, C. R., Ralph, M. A. L., and Rogers, T. T. (2023). Decoding semantic representations in mind and brain. Trends in cognitive sciences.

Giari, G., Leonardelli, E., Tao, Y., Machado, M., and Fairhall, S. L. (2020). Spatiotemporal properties of the neural representation of conceptual content for words and pictures–an meg study. Neuroimage, 219:116913.

Gramfort, A., Luessi, M., Larson, E., Engemann, D. A., Strohmeier, D., Brodbeck, C., Goj, R., Jas, M., Brooks, T., Parkkonen, L., et al. (2013). Meg and eeg data analysis with mne-python. Frontiers in neuroscience, page 267.

Groen, I. I., Dekker, T. M., Knapen, T., and Silson, E. H. (2022). Visuospatial coding as ubiquitous scaffolding for human cognition. Trends in Cognitive Sciences, 26(1):81–96.

Guggenmos, M., Sterzer, P., and Cichy, R. M. (2018). Multivariate pattern analysis for meg: A comparison of dissimilarity measures. Neuroimage, 173:434–447.

Hauk, O., Davis, M. H., Ford, M., Pulvermüller, F., and Marslen-Wilson, W. D. (2006). The time course of visual word recognition as revealed by linear regression analysis of erp data. Neuroimage, 30(4):1383–1400.

Haynes, J.-D. and Rees, G. (2006). Decoding mental states from brain activity in humans. Nature reviews neuroscience, 7(7):523–534.

Hebart, M. N., Bankson, B. B., Harel, A., Baker, C. I., and Cichy, R. M. (2018). The representational dynamics of task and object processing in humans. Elife, 7:e32816.

Humphreys, G. F., Hoffman, P., Visser, M., Binney, R. J., and Lambon Ralph, M. A. (2015). Establishing task-and modality-dependent dissociations between the semantic and default mode networks. Proceedings of the National Academy of Sciences, 112(25):7857–7862.

Hyvärinen, A. and Oja, E. (2000). Independent component analysis: algorithms and applications. Neural networks, 13(4-5):411–430.

Iamshchinina, P., Karapetian, A., Kaiser, D., and Cichy, R. M. (2022). Resolving the time course of visual and auditory object categorization. Journal of Neurophysiology, 127(6):1622–1628.

Kaiser, D., Azzalini, D. C., and Peelen, M. V. (2016). Shape-independent object category responses revealed by meg and fmri decoding. Journal of neurophysiology, 115(4):2246–2250.

King, J.-R. and Dehaene, S. (2014). Characterizing the dynamics of mental representations: the temporal generalization method. Trends in cognitive sciences, 18(4):203–210.

Konkle, T. and Caramazza, A. (2013). Tripartite organization of the ventral stream by animacy and object size. Journal of Neuroscience, 33(25):10235–10242.

Kriegeskorte, N., Mur, M., and Bandettini, P. A. (2008). Representational similarity analysis-connecting the branches of systems neuroscience. Frontiers in systems neuroscience, page 4.

Kumar, M., Federmeier, K. D., Fei-Fei, L., and Beck, D. M. (2017). Evidence for similar patterns of neural activity elicited by picture-and word-based representations of natural scenes. Neuroimage, 155:422–436.

Leonardelli, E., Fait, E., and Fairhall, S. L. (2019). Temporal dynamics of access to amodal representations of category-level conceptual information. Scientific reports, 9(1):239.

MacGregor, L. J., Pulvermüller, F., Van Casteren, M., and Shtyrov, Y. (2012). Ultra-rapid access to words in the brain. Nature communications, 3(1):711.

Maris, E. and Oostenveld, R. (2007). Nonparametric statistical testing of eeg-and meg-data. Journal of neuroscience methods, 164(1):177–190.

Miozzo, M., Pulvermüller, F., and Hauk, O. (2015). Early parallel activation of semantics and phonology in picture naming: Evidence from a multiple linear regression meg study. Cerebral Cortex, 25(10):3343–3355.

Peelen, M. and Downing, P. (2022). Testing cognitive theories using multivariate pattern analysis of neuroimaging data.

Pobric, G., Jefferies, E., and Ralph, M. A. L. (2007). Anterior temporal lobes mediate semantic representation: mimicking semantic dementia by using rtms in normal participants. Proceedings of the National Academy of Sciences, 104(50):20137–20141.

Proklova, D., Kaiser, D., and Peelen, M. V. (2016). Disentangling representations of object shape and object category in human visual cortex: The animate–inanimate distinction. Journal of cognitive neuroscience, 28(5):680–692.

Ralph, M. A. L., Jefferies, E., Patterson, K., and Rogers, T. T. (2017). The neural and computational bases of semantic cognition. Nature reviews neuroscience, 18(1):42–55.

Santi, A., Friederici, A. D., Makuuchi, M., and Grodzinsky, Y. (2015). An fmri study dissociating distance measures computed by broca’s area in movement processing: clause boundary vs. identity. Frontiers in psychology, 6:654.

Sassenhagen, J. and Draschkow, D. (2019). Cluster-based permutation tests of meg/eeg data do not establish significance of effect latency or location. Psychophysiology, 56(6):e13335.

Simanova, I., Van Gerven, M., Oostenveld, R., and Hagoort, P. (2010). Identifying object categories from event-related eeg: toward decoding of conceptual representations. PloS one, 5(12):e14465.

Singer, J. J., Cichy, R. M., and Hebart, M. N. (2023). The spatiotemporal neural dynamics of object recognition for natural images and line drawings. Journal of Neuroscience, 43(3):484–500.

Taulu, S. and Simola, J. (2006). Spatiotemporal signal space separation method for rejecting nearby interference in meg measurements. Physics in Medicine & Biology, 51(7):1759.

Tyler, L. K., Cheung, T. P., Devereux, B. J., and Clarke, A. (2013). Syntactic computations in the language network: characterizing dynamic network properties using representational similarity analysis. Frontiers in psychology, 4:271.

